# The Evolution Of Colouration And Opsins In Tarantulas

**DOI:** 10.1101/2020.04.25.061366

**Authors:** Saoirse Foley, Vinodkumar Saranathan, William H. Piel

**Affiliations:** Department of Biological Sciences, National University of Singapore, 117543, Singapore; Division of Science, Yale-NUS College, 10 College Avenue West, 138609, Singapore; Lee Kong Chian Natural History Museum, National University of Singapore, 117377, Singapore; NUS Nanoscience and Nanotechnology Institute (NUSNNI-NanoCore), National University of Singapore, 117581, Singapore

## Abstract

Tarantulas paradoxically exhibit a diverse palette of vivid colouration despite their crepuscular to nocturnal habits. The evolutionary origin and maintenance of these colours remains a mystery. In this study, we reconstructed the ancestral states of both blue and green colouration in tarantula setae, and tested how these colours correlate with the presence of stridulation, urtication, and arboreality. Green colouration has likely evolved at least eight times, and blue colouration is likely an ancestral condition that appears to be lost more frequently than gained. While our results indicate that neither colour correlates with the presence of stridulation or urtication, the evolution of green colouration appears to depend upon the presence of arboreality, suggesting that it likely originated for, and functions in, crypsis through substrate matching among leaves. We also constructed a network of opsin homologs across tarantula transcriptomes. Despite their crepuscular tendencies, tarantulas express a full suite of opsin genes – a finding that contradicts current consensus that tarantulas have poor colour vision on the basis of low opsin diversity. Overall, our results support the intriguing hypotheses that blue colouration may have ultimately evolved via sexual selection and perhaps proximately be used in mate choice or predation avoidance due to possible sex differences in mate-searching.

## INTRODUCTION

The large, hirsute spider family Theraphosidae includes some of the most visually striking spiders. Commonly known as tarantulas, these animals are notable for the fact that they are often brightly coloured, with violet, blue, green, red, and purple hues in particular, present throughout the family^1,2^. However, a long-standing enigma has been the evolutionary origin and maintenance of such vivid colouration in a group that is almost entirely crepuscular, if not nocturnal^3,4^. The evolution of colouration in tarantulas has previously received only scant attention in the literature, cf. Hsiung *et al*^2^. A well-resolved phylogeny for this group did not exist until recently^5^ and previous attempts^2^ to reconstruct the ancestral states of blue colouration were based upon a poorly resolved and outdated supertree. Hsiung, *et al*.^2^ also used vaguely defined criteria to identify blue and green colouration. For instance, no data was provided to validate their colour scorings, some of which appear to be erroneous (*E*.*g*. blue colouration was listed as present in *Haplocosmia* / *Nhandu*, and absent in *Cyrtopholis*; but see Table S3).

Here, we reconstruct the evolutionary histories of both blue (“blueness”) and green (“greenness”) integumentary colouration in the setae of tarantulas via quantitative ancestral state reconstructions. We use a robust phylogeny^5,6^ and set strict criteria for assigning colour states by calculating just-noticeable difference (JND) values^7^ for at least one representative species for each genus represented in our phylogeny. In order to address the function of colours in tarantulas, we also constructed data matrices based on prior publications detailing the presence or absence of (i) stridulatory setae, (ii) urticating setae, and (iii) arboreal tendencies for each genus in this study, and conducted a series of correlative tests to determine whether any of these traits might inform the function of blue or green colouration in tarantulas. We hypothesize that blue colour should be associated with stridulating / urticating setae, or both, if it either originally evolved or has been exapted (sensu^8^) for an aposematic function. However, we posit that green colouration could aid in crypsis and should show an association with arboreality.

It is generally thought that tarantulas possess little or no colour vision, with only a weak colour spectrum discrimination^2,9^, with Hsiung, *et al*.^2^ noting that sexual selection as a driving force behind the evolution of tarantula colouration is unlikely. Morehouse *et al*.^10^ argue that tarantulas are colour-blind on the basis of finding a low opsin diversity in *Aphonopelma*^10^. We investigate whether this claim holds true for other tarantulas by scoring the presence and relative expression of opsins from transcriptomic data in 25 tarantula genera. Opsins are specialized, transmembrane proteins belonging to the superfamily of G-protein coupled receptors that convert light photons to electrochemical signals through a signalling cascade^11^. While the expression of multiple opsin species does not necessarily imply greater colour discrimination, the absence of opsin orthologs could provide support to the hypothesis that tarantulas, broadly speaking, cannot perceive colours.

## RESULTS

### Phylogenetic Signal of Colour Traits

Our phylogeny contains 37 theraphosid genera and 2 outgroups (*Damarchus* sp. and *Linothele* sp.), covering the entire spectrum of theraphosid subfamilies minus Selenogyrinae. Blueness occurred in 25 genera and both outgroups, greenness in 11 genera, stridulation in 25, urtication in 14, and arboreality in 9 (Table S3). Blomberg’s K and Pagel’s λ values are reported in Table 1. Only weak phylogenetic signal was detected in each case, with K’s ranging from 0.38-0.46, and λ’s from 0-0.42. Only 4 internal nodes in the ancestral state reconstructions of blue had posterior probabilities of ≥95%, and no such nodes were recovered in either the green and arboreality reconstructions (Figures S1, S2, and S3).

**Table 1:**
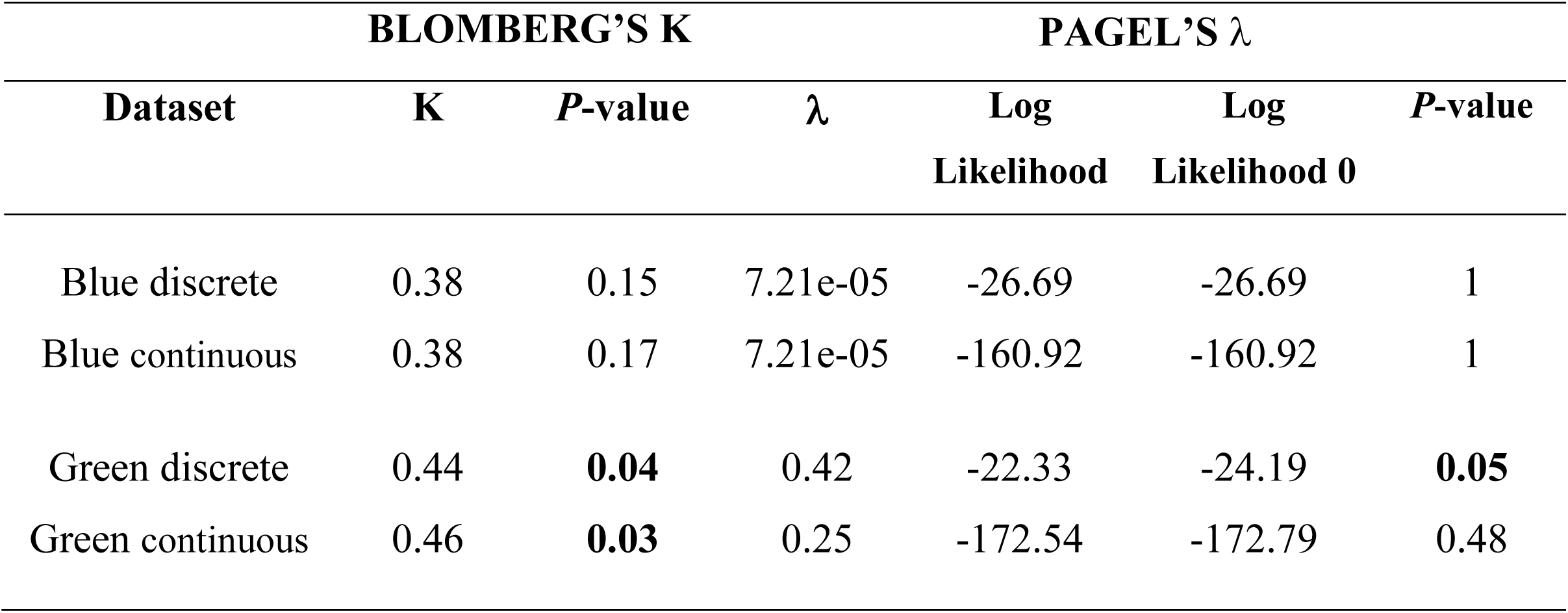
Phylogenetic signal as determined by Blomberg’s K and Pagel’s λ. Blomberg’s K values < 1 inform that there is less phylogenetic signal than would be expected under Brownian motion. Pagel’s λ informs whether the phylogeny alone is sufficient or not to explain the distribution of traits.

Trait correlation outputs for all traits and colours are reported in Table 2. The only association that displayed a significant *P*-value is “greenness depending on arboreality” (0.01). This informed the need to subsequently conduct dependence tests.

**Table 2:** Correlative tests as derived via Pagel’s test. *P*-values indicate that blueness does not appear to have any correlation with any traits tested, and greenness is only correlated when a model where it depends upon arboreality is selected.

| COLOUR | Trait | Likelihood Ratio | *P*-value |
| --- | --- | --- | --- |
| **BLUE** | Stridulation | 2.82 | 0.59 |
|  | Urtication | 1.02 | 0.90 |
|  | Arboreality | 3.42 | 0.49 |
|  | Depends on Arboreality | 3.42 | 0.18 |
| **GREEN** | Stridulation | 4.0 | 0.40 |
|  | Urtication | 9.09 | 0.06 |
|  | Arboreality | 9.14 | 0.06 |
|  | Depends on Arboreality | 8.69 | **0.01** |

### Evolution of Blueness

The ancestral state reconstruction of continuous blue (ΔE JND values), with ancestral nodes annotated according to the reconstruction of discrete blue data, is shown in Figure 1. This shows that blue was likely present at the root of the tree, but was subsequently lost and regained multiple times. Of the five gains, three are at terminal nodes in Old World subfamilies, one is relatively shallow (the ancestor to *Orphnaecus* and *Phlogiellus* also in the Old World, and one is at the root of our phylogeny and is largely retained throughout the subfamilies Ornithoctoninae, Poecilotherinae, Harpactirinae, Theraphosinae, Psalmopoeinae, Aviculariinae, and some Ischnocolinae. Blueness is also reported in both of our non-tarantula outgroups (*Linothele* and *Damarchus*), and only two of the included theraphosid subfamilies lack any blue members (Schismatothelinae and Stromatopelminae). Blueness appears to have been lost seven times, with five of those losses occurring at the tips or relatively shallow nodes, especially in Neotropical subfamilies. A full *ancThresh* reconstruction for blueness, with posterior probabilities as pie charts plotted at every node, is reported in Figure S1.

**Figure 1:**
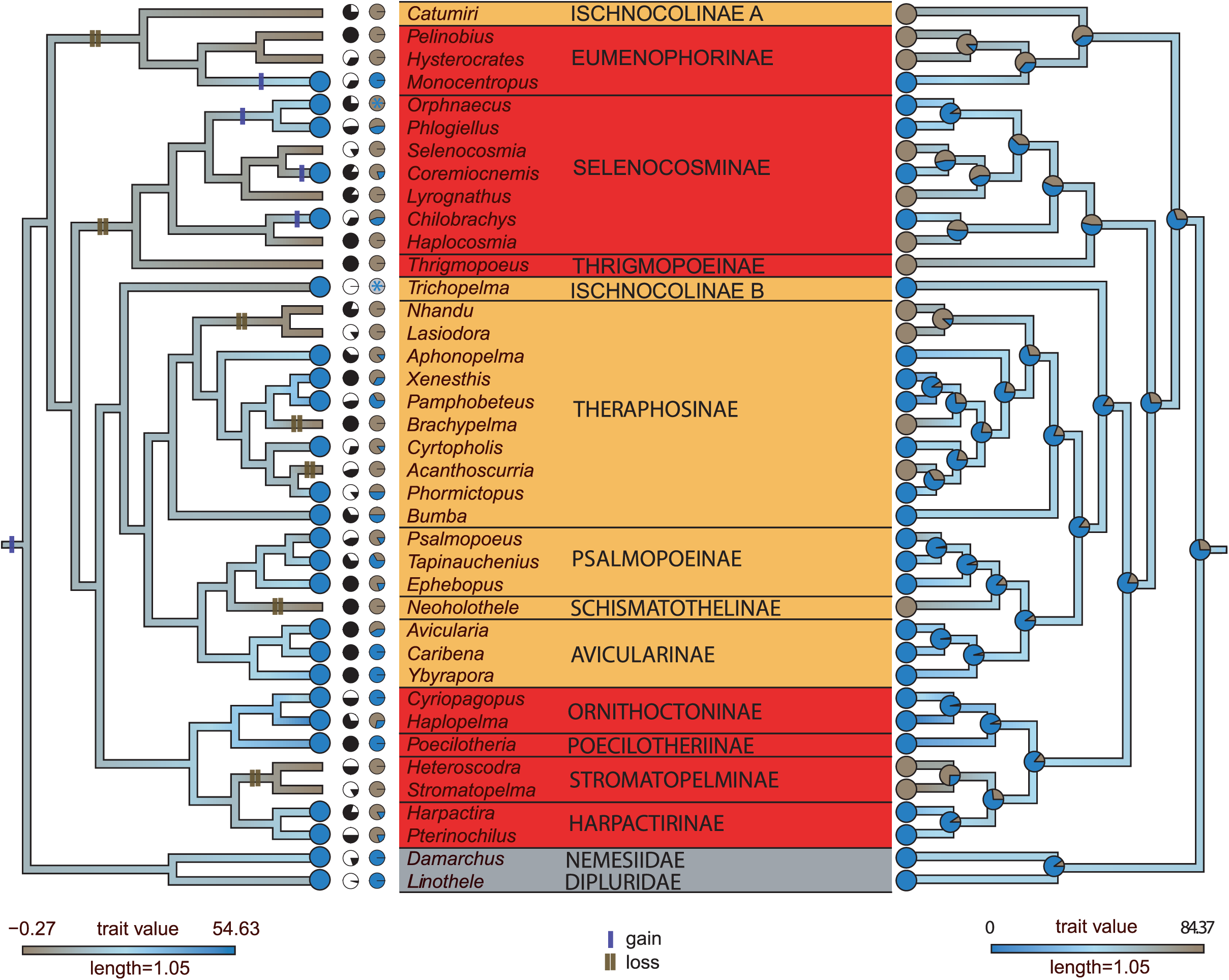
Blueness mapped onto our phylogeny^5,6^ as inferred from the ancestral state reconstructions of the continuous ΔE *Lab* values (Right), with the posterior probabilities of ancestral nodes plotted as pie charts. Left panel shows our conservative ΔE *b* mapping, with the discretization of the ΔE *Lab* values (Right) suggesting 5 gains (single blue bars) and 7 losses (double brown bars). Old World subfamilies are shown in red, and New World subfamilies in orange. Pie charts adjacent to taxon names depict the proportions of currently described species in each genus with digital photographs available (black = images found, white = no images), and the proportions of species with images that possess blue colouration (grey = no available images) – see Table S7. Blue * denote undescribed species that can be unambiguously attributed to the corresponding genera with blue colour.

### Relationship Between Greenness and Arboreality

Both raw and weighted AIC values to assess the relationships between greenness and arboreality are reported in Table 3. A mirror tree showing ancestral state reconstruction of continuous green on the left, and arboreality on the right, is shown in in Figure 2. Ancestral nodes were annotated according to the reconstruction of discrete green data with *ancThresh*. Both arboreality and greenness is reported in six genera, but a further eight genera possess either greenness or arboreality. Greenness does not appear to have been lost at any point throughout the evolution of tarantulas, but has been gained at least eight times. Six of these eight gains appear at terminal nodes (tips). Arboreality shows five independent gains, of which two occur at the tips. Full *ancThresh* reconstructions with posterior probabilities and pie charts plotted at every node for arboreality and greenness are reported in Figures S2 and S3 respectively.

**Table 3:** AICs and weighted AICs (relative probabilities) to assess four different evolutionary scenarios for greenness, where models of different evolutionary relationships are tested in each case. A scenario where greenness depends on the prior evolution of arboreality is supported (AIC - 86.64, weighted AIC - 0.5).

| DEPENDENCY | AICs | WEIGHTED AICs |
| --- | --- | --- |
| Independent | 81.47 | 0.03 |
| Arboreality depends on colour | 77.77 | 0.19 |
| Colour depends on arboreality | **75.13** | **0.72** |
| Interdependent | 80.32 | 0.054 |

**Figure 2:**
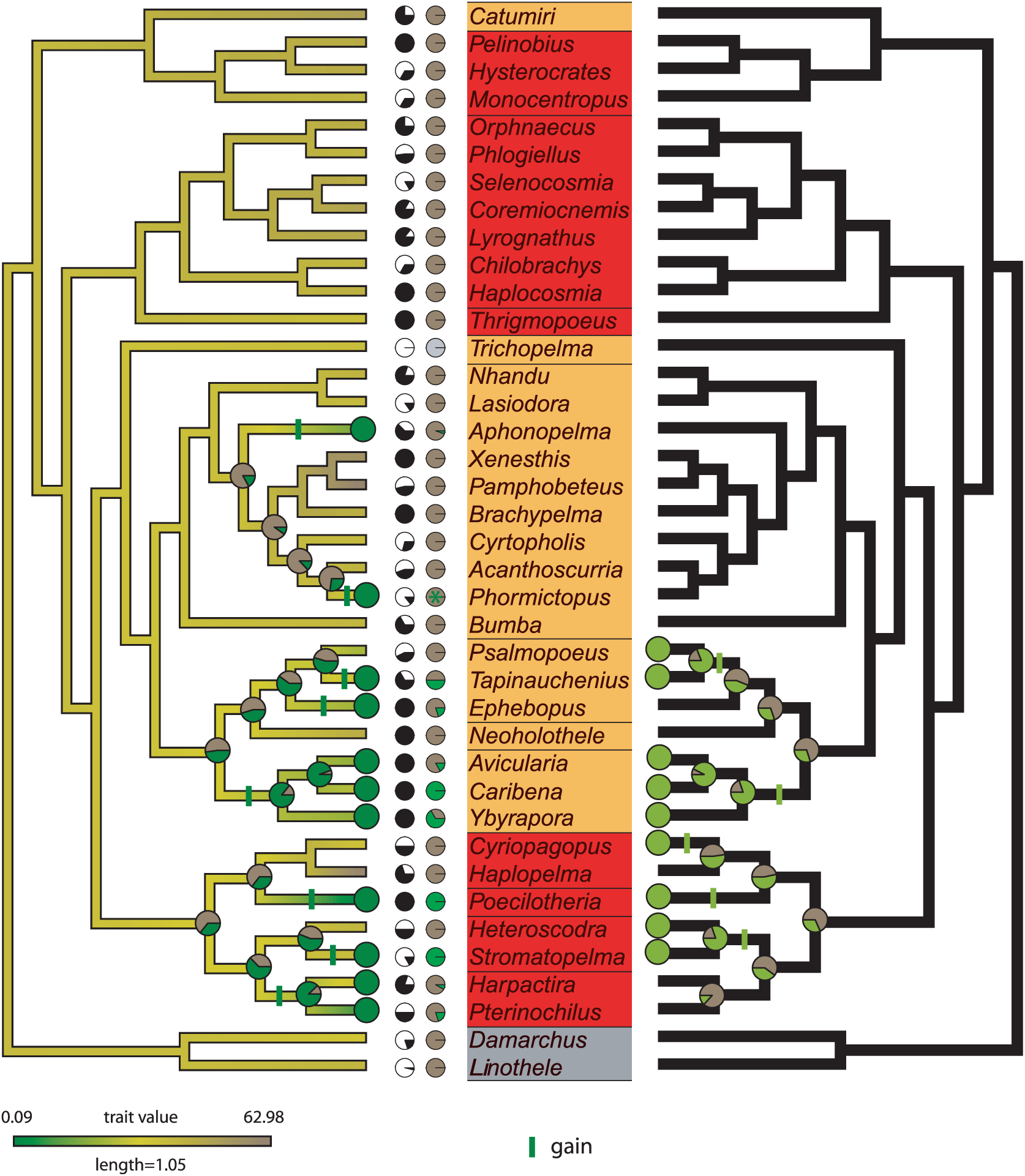
Mirror topologies showing the distribution of greenness (left) and arboreality (right). Posterior probabilities for select (for clarity) ancestral nodes are shown as pie charts. Trait gains are represented by single green bars. In the left panel, trait values and branch shading refer to the ancestral state reconstructions of continuous ΔE *a* values. The ancestral posterior probabilities of continuous ΔE *Lab* values of relevant nodes that were used to discretize the greenness trait are also shown. Greenness is gained eight times, primarily at the tips with no loss. Arboreality is gained five times, and is never lost. Pie charts adjacent to taxon names depict the proportions of currently described species in each genus with digital photographs available (black = images found, white = no images), and the proportions of species with images that possess green colouration (grey = no available images) – see Table S7. Green * denote undescribed species that can be unambiguously attributed to the corresponding genera with green colour.

### Opsin Diversity in Tarantulas

Our opsin homolog network is presented in Figure 3. There are 192 nodes total, with each taxon for which data was available being represented by at least one sequence. While homologs were recovered from some transcriptomes generated from leg-only samples, transcriptomes generated from whole body samples were most successful for recovering the full suite of opsin homologs. Generally, the degree (number of connected edges) of all reference opsins was quite high, with SWS = 57, MWS = 65, PER = 69, RH1 = 80, RH2B = 91, and RH2A = 107. Many nodes demonstrate homology to several opsin types, with some demonstrating homology to all reference opsins. Complete opsin homolog and ortholog data for all species is available in Table S6.

**Figure 3:**
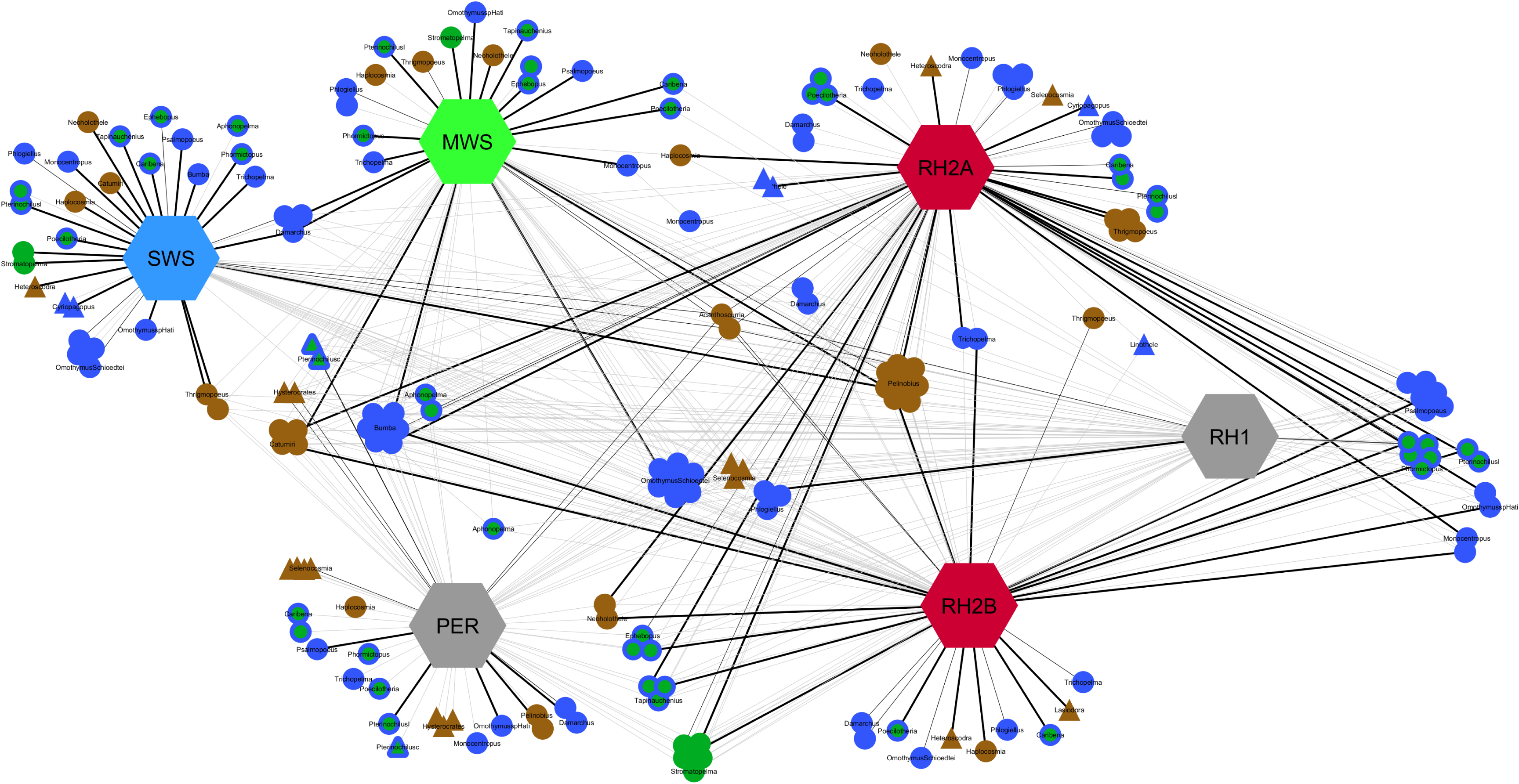
Network of opsin sequences in tarantulas. Thick, black edges are cases where all three tools predict the homology, and the connected nodes can be considered orthologous. Thin, black edges connect nodes when any two tools agree on the homology. Grey edges connect nodes when only a single tool agrees on the homology. Reference opsins are arranged in increasing order of wavelength sensitivities. The presence of blue and / or green colour in each genus is indicated. Reference opsins are represented by hexagonal nodes, whole body samples by circular, and leg-only samples by triangles.

The phylogenetic relationships among opsin families in tarantulas is shown in Figure S4. It is largely consistent with results from other invertebrate studies^12^, but differs in that peropsins (RRH) emerge as the most derived in our case, where they emerge as the most basal in Biscontin, *et al*.^12^. Full sequence alignments (MSA) of tarantula opsins are available in Figure S5. Generally, even at positions where individual amino acids vary in each MSA, they demonstrate functional conservation.

## DISCUSSION

### Blueness is Likely Ancestral to Theraphosidae

As seen in Figure 1, the outgroup taxa are blue. This polarity, along with the posterior probabilities on the ancestral-most nodes, suggest that the earliest tarantulas were likely blue. This early origin of blueness appears to be retained in the present-day members of the New World subfamilies Theraphosinae, Aviculariinae, and Psalmopoeinae (and at least some of the taxonomically unresolved Ischnocolinae), with 4 losses distributed mostly at the tips or shallower nodes. Blueness is even retained throughout some Old World subfamilies, namely Ornithoctoninae, Poecilotheriinae, and Harpactirinae, while it has been lost twice in older clades and re-evolved four times at the tips or shallower nodes.

While Hsiung, *et al*.^2^ also recovered blueness as ancestral, our estimates of the frequency and pattern of evolution differ. Hsiung, *et al*.^2^ suggested that blueness evolved at least eight independent times in tarantulas, and that the true number of gains could be much greater still. Hsiung, *et al*.^2^ supported their claim that blueness evolved many times independently in part by arguing that distinctly different nanostructures have converged to produce the same colour. By comparison, our results (Figure 1) indicate that blueness has evolved at most five times, and was lost more often than it was gained (seven times). Interestingly, of the five gains reported here, all occur in Old World groups (three tips), and five of our seven losses are in New World groups at the tips or relatively shallow nodes.

Hsiung, *et al*.^2^ also characterised the photonic nanostructures within the setae of selected blue theraphosid tarantulas as being either amorphous spongy or multi-layer, and concluded that these distinct nanostructures are a product of convergent evolution to produce similar blue colours. They argued against sexual selection as the driver behind blue colouration because typically sexual selection causes colour divergence, not convergence, at least in sympatry^13,14^. However, their character scorings are based on apparently low-resolution and low-magnification 2D transmission electron microscopy (TEM) images of what are otherwise complex nanostructures. The EM data for species with multilayer nanostructures in Hsiung, *et al*.^2^ suggests that the nanostructure can be more disordered, especially toward the setal interior. This is analogous to the interplay between order and disorder often seen in the same individual, for instance in many beetles^15,16^. In addition, some amount of disorder could be an artefact of the sectioning and invasive TEM imaging process. The TEM data in Hsiung, *et al*.^2^ together with precise, bulk 3D SAXS structural information from taxa in both Old World (Ornithoctoninae and Poecilotheriinae) and New World (Psalmopoeinae) theraphosid subfamilies^16^ suggests there is only one fundamental type of setae photonic nanostructure in tarantulas — a perforated lamellar or a multilayer nanostructure with random perforations as seen in certain species of Polyommatine and Papilionid butterflies^17,18^, with differing degrees of structural disorder.

These observations are consistent with a single ancestral origin of blue colouration in tarantulas and thus does not preclude a role for sexual selection in their evolution. It is well appreciated that certain tarantula species exhibit sexual dimorphism, including age-related dichromatism across several genera e.g. *Haplopelma, Xenesthis*^*2*,19,^ *Psalmopoeus*^*20*,^ *Aphonopelma*^*21*,^ and many Aviculariinae^22^. Penultimate males frequently display vivid metallic colourations, and having examined a mature *Pamphobeteus antinous* male with vivid iridescent blue bristle ornamentation on the legs and pedipalps, Simonis, *et al*.^23^ proposed that such colouration might play a role in sexual signalling.

Alternatively, given that upon maturity many male tarantulas will leave their retreats and wander in search of females^24-26^, it is plausible that different selective pressures other than sexual selection could be driving sexual dichromatism. For instance, certain colours are either aposematic or cryptic under moonlight^27,28^, and therefore useful to the wandering male but not to the fossorial female.

### Does Arboreality Precede the Evolution of Greenness?

Blomberg’s K values for greenness (Table 1) support the hypothesis that green taxa tend to cluster together in our phylogeny. Despite the small K values, the significant *P*-values imply that closely related taxa are probably more similar in their greenness than random pairs chosen from across the phylogeny. However, Pagel’s λ analyses indicate that the phylogeny alone cannot explain the observed patterns. When both measures are considered together, it’s reasonable to infer that it is slightly more probable to find greenness clustered among closely related taxa, but the overall structure of the topology fails to explain why this is the case. This observation is generally consistent with Figure 2. One can observe that if a given genus exhibits either arboreality or greenness, then its close relatives also tend to exhibit one or both traits.

In agreement with Hsiung, *et al*.^2^, green colouration was found to be rare in our phylogeny. However, our ancestral state reconstruction (Figure 2) has revealed a previously undocumented relationship between greenness and arboreality. Of the 11 green genera in this study, six taxa are arboreal. Both traits appear to have evolved within the genus of *Poecilotheria* (*i*.*e*. emerged as tip states in this analysis). For *Avicularia, Caribena*, and *Ybyrapora*, both greenness and arboreality are shown to emerge in the common ancestor to the three. However, in the cases of *Stromatopelma* and *Tapinauchenius*, arboreality is reported in the immediate ancestors of these genera, but greenness is only reported as a tip-state in each – arboreality preceded the evolution of greenness. Our dependency tests (Table 3) support an evolutionary scenario where evolving greenness depends on the *a priori* presence of arboreality. We hypothesize that this is the case for *Stromatopelma* and *Tapinauchenius*, with some caveats (see below). We cannot conclusively say whether this is true for either *Poecilotheria* or the subfamily Aviculariinae (*Avicularia, Caribena*, and *Ybyrapora*), but the addition of further taxa at the species level could assist in determining whether greenness or arboreality emerged first in these clades.

There are five genera for which greenness is reported without arboreality. Of those five, neither *Aphonopelma* nor *Phormictopus* have any close arboreal relatives. However, *Ephebopus, Pterinochilus*, and *Harpactira* are all sister to arboreal clades that do contain green members. The case of *Ephebopus* is particularly interesting. It is the only fossorial member of the subfamily Psalmopoeinae, which includes the above-mentioned *Tapinauchenius*, and greenness is only restricted to a small abdominal patch in our representative species (*E. cyanognathus*). West, *et al*.^29^ reported that certain *Ephebopus* species may colonize arboreal domains, e.g. early instars of *E. murinus* have been observed creating tube webs in bromeliads, and *E. rufescens* can be found in hollow rocks and tree-holes. The study also noted, however, that *Ephebopus* is primarily a fossorial genus, and even those species that display arboreal tendencies are more often found in burrows, which informed our treatment of the genus as non-arboreal. Within the context of our results, it is plausible to speculate that, as a genus whose members contain localized patches of green along with fossorial species that occasionally display arboreal tendencies, *Ephebopus* may represent a taxon that is in the process of evolving an increasingly arboreal lifestyle, concomitant with the emergence of greenness and the crypsis that it confers.

Four genera are listed as arboreal and non-green. *Heteroscodra*’s closest relative, *Stromatopelma*, is both arboreal and green, and the sister clade to the pair also contains green members (*Pterinochilus* and *Harpactira*). *Cyriopagopus* forms a clade with *Poecilotheria* (a genus that’s both arboreal and green) and *Haplopelma* (fossorial and non-green). For *Psalmopoeus*, the ΔE *b* value for this genus was calculated to be 53 (Table S3), which was very close to the cut-off (<50) required to be considered green. Its closest relatives in this study all contain green members.

### Opsin Diversity and Evolution in Tarantulas

Our results document that the same suite of opsins as in jumping spiders are likewise expressed in a wide array of tarantula species. In particular, both homologs and orthologs to short-wavelength and medium-wavelength sensitive opsins that can detect blues and greens^30^ were found in all whole-body transcriptomes, except for *Hysterocrates* sp., *Lasiodora parahybana, Pterinochilus chordatus*, and the diplurid *Linothele* sp. Morehouse, *et al*.^10^ reported that *Aphonopelma* only possesses RH2A and UV-sensitive short-wavelength opsins. While we acknowledge that some of our tarantula opsin sequences show homology to many types of reference opsins, our results nonetheless suggest that Morehouse *et al*.’s observation underestimates true opsin diversity among tarantulas when more broadly sampled. All subfamilies possess the full complement of typical arthropod opsins. Given that jumping spiders are known to discriminate colours, including reds, greens, and blues^30^, we cannot discount the possibility that tarantulas are able to perceive colours based on our results (Figure 3). While Dahl and Granda ^9^ noted that *Aphonopelma chalcodes* lack the neurological response complexities seen in colour discriminating eyes, we encourage future behavioural studies from a variety of genera to see if this reported inability is broadly representative of tarantulas. In short, we cannot rule out that blue colouration could not have evolved in tarantulas for the purposes of sexual signalling due to their poor eyesight e.g.^2^, or on the basis of low opsin diversity. However, given that blueness is lost more frequently than it is gained, we suggest that it is perhaps under weak selection or has been exapted for cryspis or aposematic signalling in twilight or perhaps moonlight^27,28^.

## CONCLUSIONS

Our research offers novel insights into the evolutionary origin and potential functions of blue and green colouration in tarantulas. We report that blueness is an ancestral condition for tarantulas and was likely present among the groups’ earliest representatives. This finding leaves open the possibility that blueness may have originally evolved for the purposes of sexual signalling, and may continue to serve in this role given that most tarantulas still possess the typical, full suite of arthropod opsins. On the other hand, greenness has likely evolved at least eight times, primarily at the tips, and does not appear to have been lost at any point in the evolution of tarantulas. We provide support for a conditional relationship where evolving greenness depends likely on the prior presence of arboreality, and compare the ancestral state reconstructions of both greenness and arboreality to further support this idea. We hope that future work will include comparative studies on the visual perception and putative mate-choice experiments in tarantulas to better understand the selective pressures involved in the origin and maintenance of vivid colouration in this enigmatic group.

## DATA ACCESSIBILITY

All data used for this analysis can be found in this article and in the electronic supplementary material.

## AUTHOR CONTRIBUTIONS

SF and VS designed the study. SF compiled the data matrices. VS and SF coded and performed the phylogenetic analyses of coloration in R. SF generated the transcriptomic results with input from VS and WHP. SF and VS analysed the results. SF and VS wrote the manuscript with input from WHP. The authors declare no competing interests, financial or otherwise.

## ACKNOWLEDGMENTS

We thank Nate Morehouse and Megan Porter for kindly sharing their tarantula opsin sequence data. The computational results reported in this study were possible thanks to a Yale-NUS College Shared Instrumentation Grant (IG15-SI101) to VS and WHP. VS acknowledges support from Yale-NUS start-up funds. WHP acknowledges support from the South East Asian Biodiversity Genomics (SEABIG) centre, the Singapore Ministry of Education (grant number MOE2016-T2-2-137), and by Yale-NUS College (grant numbers IG14-SI002 and R-607-265-052-121).

## METHODS

### Phylogeny Construction

The phylogeny published by Foley, *et al*.^5^, based on 2460 orthologous genes was used as a constraint tree for this study, but taxa were reduced to the genus level for ease of assigning character states in subsequent steps. The R package “PASTIS”^34^ facilitated the inclusion of a further 11 genera that lack sequence data by constraining their clade memberships as determined by previous studies (Table S1). Branch lengths for these additional taxa were estimated by MrBayes v3.2^35^, which ran under a birth-death model for 100,000 generations, with sampling every 160 generations and the burnin set to 25%.

### Colour Survey

Photographs corresponding to representative species of each genus in the phylogeny were primarily acquired from arachnologist Rick C. West’s photographic archive (http://birdspiders.com), but other sources included Gaia Biological (http://thespidershop.co.uk), Nicky Bay (http://nickybay.com), and the published literature. A list of representative species per genus, as well as the source for each photograph used in this study, can be found in Table S2. These photographs were analysed with ImageJ v1.52.

Each photo was individually examined with the “Colour Inspector 3D” ImageJ plugin and visualized in the CIELAB colour space. The “bluest” regions of each photo were characterized by greater negative *b* values, and the “greenest” regions by greater negative *a* values. These bluest and greenest regions of each photo were then cropped and reloaded into ImageJ as new images. The “Colour Transformer” plugin was used to convert each of these images into a stacked CIELAB image. The “Measure” command was used on each frame to obtain mean *L, a*, and *b* values for each taxon, as well as for six reference leaf images. The *Lab* values for the six leaves were averaged and used as reference green colour values, and the *Lab* values for the taxon with the greatest -*b* value (i.e. the “bluest” species, *Haplopelma lividum*) was chosen to be our reference for blue values. We also obtained *Lab* values for the jumping spider, *Maratus splendens*, as a potential blue reference^36^. However, their *-b* values were less extreme than our chosen reference, *Haplopelma lividum*.

Using the conservative International Commission on Illumination (CIE) metric for colour discrimination, ΔE 76^37^, four separate Euclidean difference values were then calculated for each taxon in order to quantify the perceived colour difference of each taxon relative to the reference colour values: 1. ΔE *Lab* against the blue reference, 2. Δ*b* against the blue reference, 3. ΔE *Lab* against the green reference, and 4. Δ*a* against the green reference. Values 1 and 3 informed our continuous colour dataset, whereas values 2 and 4 were discretized, with any taxon scoring <50^37^ being scored as possessing the corresponding colour (i.e. “1”; for values < 50 indicate colours are more similar than opposite). Values 2 and 4 also served to correct for varying light intensities across images, as only the actual colour values were considered in these cases without the influence of *L* affecting further calculations. Other methods were considered for colour discrimination, but they were ultimately deemed too permissive. For example, the same discriminatory criteria under ΔE 2000^38^ would result in all but a single species in our phylogeny being considered blue. All measurements and instances where our colour scoring differs from Hsiung, *et al*.^2^ are noted in Table S3.

### Habitat and Defensive Trait Survey

A discrete data matrix was constructed based on the presence or absence of the following traits: Urticating setae were marked as present in genera from the subfamilies Theraphosinae^39^ and Aviculariinae^22^, as well as the genus *Ephebopus*^40,41^. Stridulatory setae were marked as present in *Chilobrachys, Coremiocnemis, Haplocosmia, Lyrognathus, Orphnaecus, Phlogiellus, Poecilotheria, Selenocosmia*^42^, *Acanthoscurria, Brachypelma, Cyrtopholis, Lasiodora, Nhandu, Pamphobeteus, Phormictopus*^43^, *Cyriopagopus, Haplopelma, Harpactira, Hysterocrates, Monocentropus, Pelinobius, Psalmopoeus*^44^, *Bumba*^45^, *Pterinochilus*^46^, and *Thrigmopoeus*^47^. Lastly, arboreal tendencies were marked as present in *Avicularia, Caribena, Heteroscodra, Poecilotheria, Psalmopoeus, Stromatopelma, Tapinauchenius*^48^, *Cyriopagopus*^49^, and *Ybyrapora* because its members have historically been treated within *Avicularia*. All data is available in Table S3.

### Ancestral State Reconstructions and Correlative Tests

Phylogenetic signal of discrete blue, discrete green, Δ*b* values against the blue reference, and Δ*a* values against the green reference were assessed using both Blomberg’s K^50^ and Pagel’s λ^51^. A set of functions in the R package “phytools”^52^ facilitated ancestral state reconstructions and correlative tests. The ancestral states for discrete blue, discrete green, and arboreality were separately estimated using 10 million generations of Bayesian MCMC with the *ancThresh()* function^53^, and the posterior probabilities were plotted at ancestral nodes as pie charts. We coded the traits as present at any internal nodes whose corresponding posterior probabilities were ≥66% (e.g.^54,55^), and made note of any nodes whose corresponding posterior probabilities were ≥95%. The JND (continuous) colour data was then traced through the phylogeny using the *contMap()* function. The *ancThresh* pie charts corresponding to ancestral nodes whose posterior probabilities indicated that a trait was present were subsequently overlaid onto the corresponding *contMap* outputs for ease of interpretation.

Pagel’s^56^ models of correlated evolution were applied to our discrete data to estimate the relationships between the presence of blue or green and habitat and defensive traits using the *fitPagel()* function. Additionally, sets of AIC values were generated with *fitPagel* to estimate more specific relationships between colours and arboreality, *i*.*e*. whether either trait depends on the other, or whether the traits are independent / interdependent. The *fitDiscrete()* function was used in each case.

### Generating an Opsin Homolog Network

We assembled publicly available tarantula transcriptomes for each genus in our phylogeny that had data available (data sources in Table S4). We then acquired candidate opsin sequences from six different types of opsin proteins that other spiders are known to possess according to^10^: middle-wavelength-sensitive rhodopsin-like-1, 2A and 2B (RH1, RH2A and RH2B respectively), medium-wave-sensitive opsin-1 (MWS), short-wave-sensitive opsin-1 (SWS), and retinal pigment epithelial RPE-specific rhodopsin (RRH). Opsin accession numbers are available in Table S5. These six representative sequences served as queries for BLASTP^57^ searches against our transcriptome set. Only hits with e-values greater than 10^−7^ were discarded, with a view towards treating these hits as a tentative list of tarantula opsin homologs.

Homology was inferred between our candidate opsin sequences and our reference opsin set for each species using three different tools: SwiftOrtho^59^, FastOrtho (http://enews.patricbrc.org/), and OrthoFinder^60^. SwiftOrtho and FastOrtho were run with e-values of 10^−40^ for homology searching to reduce potential false positives while OrthoFinder was run using default settings. Per the “moderate” criteria from the Alliance of Genome Resources^61^, hits may be considered orthologous if three or more of the twelve tools in their suite converge upon that result. Hence, any hits which are recovered by all of the above-mentioned tools can be considered orthologous. Results from each of these programs were pooled, and loaded into Cytoscape^62^ for network visualization.

The reference opsins, along with sequences corresponding to nodes that only showed a degree of 1 and were considered orthologous were then aligned using MAFFT^63^. This was to ensure that only nodes showing unambiguous orthology to a single reference opsin would be retained. alignment served as an input for RAxML v8.2.11^64^, which generated a maximum likelihood tree.

