## Supplementary Data for "The Evolution Of Colouration And Opsins In Tarantulas"

**Supplementary Data for Foley *et al.* – The Evolution of Colouration and Opsins in Tarantulas**

*Notes on Trait Coding, Previous Studies and Future Perspectives*

To best preserve the uniformity (e.g. illumination) of our colour measurements, we have made every effort to work from a single source for all photos (supplementary Table S2), where possible. We hence used only the Δ*a* and Δ*b* parameters in the absence of the *L* to inform our discrete colour datasets when assigning colour states. We endeavoured to only measure colour in species which have been formally described, but it was occasionally necessary to measure an undescribed species identifiable only to the genus level due to a lack of photographic images. We also found that probabilities of ≥95% were quite rare in all cases, and while it was too strict to be considered here, our threshold of 66% is still quite conservative in comparison to other recent studies (Kin *et al*., 2016; Boyle & Herrel, 2018). It's important to note that there are some reports of blueness occurring in certain taxa which we have marked conservatively as non-blue in this study. For instance, some mature males of *Acanthoscurria* spp. are thought to exhibit pinkish or bluish iridescence (Rogério Bertani, pers. comm.). This would certainly make sense in the context of our study — a blue member of *Acanthoscurria* would result in one less loss of blueness, and *Phormictopus* would then be part of that same ancestral gain of blueness along with the rest of the Neotropical genera. However, we opted not to consider such genera as blue due to a lack of corroborating photo material. Another important point to consider is that certain taxa are known to possess iridescent claw tufts on the undersides of their feet, e.g. *Kankuamo* spp. (Perafán *et al*., 2016). We managed to source photos for two species from which this iridescence is known — *Poecilotheria regalis* and *Pterinochilus murinus*. The specimens in the photos had their forelegs raised, and the underside could hence be measured. Unfortunately, there are no formal reports discussing the prevalence of iridescent claw tufts in tarantulas. Even though iridescent claw tufts do not appear to be of taxonomic significance, we encourage future morphological studies to nonetheless record the presence or absence of this trait in the species being examined.

The cases of *Acanthoscurria* and *Pterinochilus* also serve as potential explanations for the significant differences between our results and those from Hsiung *et al*. (2015), who considered some of our non-blue taxa to be blue (*Acanthoscurria*, *Haplocosmia, Lyrognathus, Nhandu,* and *Selenocosmia*) without providing any corresponding photos or quantitative measures. As discussed, some undescribed species might possess blue colours, but we were unable to find images or descriptions of any blue representatives from any of these genera. Hsiung *et al*. (2015) also considered *Cyrtopholis* and *Pterinochilus* to be non-blue, while we have assigned them as blue. As discussed, *Pterinochilus* possesses iridescent blue claw tufts, and our photo material corresponding to a male *Cyrtopholis gibbosa* met our requirements to be considered blue with a ΔE *Lab* value of 44.

The phylogenetic framework used in this study might also explain these differences from Hsiung *et al*. (2015), as it received strong bootstrap support (Foley *et al*., 2019). By contrast, Hsiung *et al*. (2015) used a supertree constructed from a variety of previously published studies (Bertani, Nagahama & Fukushima, 2011; West, Nunn & Hogg, 2012; Bertani, 2012; Bertani & Guadanucci, 2013; Guadanucci, 2014), and included some relationships now considered erroneous, particularly where subfamilies are concerned.

Finally, our study included just 37 of the 147 described tarantula genera at the time of writing (Kambas, 2020) — just over 25%. Robust phylogenies are key to accurately inferring ancestral states, so we chose to be quite conservative when adding taxa without sequence data in order to preserve as much information as possible in our results. The Foley *et al*. (2019) phylogeny was a sufficiently robust starting point, and we chose to only add genera whose relationships could be confidently and securely placed within that framework, as determined by a variety of other studies (West, Nunn & Hogg, 2012; Lüddecke *et al*., 2018; Hüsser, 2018; Galleti-Lima & Guadanucci, 2018; supplementary Table S1). We have confidence that future studies will indeed have access to a more complete tarantula phylogenomic and transcriptomic data to work with and ultimately illuminate and clarify the selective pressures involved in the origin and maintenance of vivid colouration.

**Table S1:** A list of taxa added to the backbone phylogeny (Foley *et al*. 2019) using "PASTIS".

| **GENUS** | **POSITION** | **REFERENCE** |
|---|---|---|
| *Avicularia* | Sister to *Caribena* | Hüsser 2018 |
| *Brachypelma* | Forms a missing clade sister to other Theraphosinae | Galleti-Lima and Guadanucci 2018 |
| *Chilobrachys* | Sister to *Haplocosmia* | West *et al*. 2012 |
| *Coremiocnemis* | Sister to *Selenocosmia* | West *et al*. 2012 |
| *Harpactira* | Sister to *Pterinochilus* | Lüddecke *et al*. 2018 |
| *Lyrognathus* | Sister to *Selenocosmia* + *Coremiocnemis* | West *et al*. 2012 |
| *Nhandu* | Sister to *Aphonopelma* | Galleti-Lima and Guadanucci 2018 |
| *Orphnaecus* | Sister to *Phlogiellus* | West *et al*. 2012 |
| *Pamphobeteus* | Forms a missing clade sister to other Theraphosinae | Galleti-Lima and Guadanucci 2018 |
| *Xenesthis* | Forms a missing clade sister to other Theraphosinae | Galleti-Lima and Guadanucci 2018 |
| *Ybyrapora* | Sister to remaining Aviculariinae | Hüsser 2018 |

**Table S2:** Links to photos used for this study. For photo pairs, the first was used to measure blueness, and the second to measure greenness. BS = Birdspiders.com.

| Genus | Species | Photo Sources |
|---|---|---|
| *Acanthoscurria* | *insubtilis* | BS |
| *Aphonopelma* | *mooreae* x2 | BS, and Fearnottarantulas.com |
| *Avicularia* | *avicularia* and *variegata* | BS, and Fukushima & Bertani 2017, figure 68 |
| *Brachypelma* | *Emilia* | BS |
| *Bumba* | *pulcherrimaklaasi* | BS |
| *Caribena* | *versicolor* x2 | Thespidershop.co.uk |
| *Catumiri* | *petropolium* | BS |
| *Chilobrachys* | *bicolor* | BS |
| *Coremiocnemis* | *hoggi* | BS |
| *Cyriopagopus* | *thorelli and sp.* "Hati Hati" | BS (under *Omothymus*), and Thespidershop.co.uk |
| *Cyrtopholis* | *gibbosa* | BS |
| *Damarchus* | *workmani* | flickr.com/photos/nickadel/ (Nicky Bay) |
| *Ephebopus* | *cyanognathus* | BS and Thespidershop.co.uk |
| *Haplocosmia* | *himalayana* | BS |
| *Haplopelma* | *lividum* | BS (under *Cyriopagopus livius*) |
| *Harpactira* | *pulchripes* and *sp.*1 RSA | BS |
| *Heteroscodra* | *maculata* | BS |
| *Hysterocrates* | *crassipes* | BS |
| *Lasiodora* | *parahybana* | BS |
| *Linothele* | *paulistana* | BS |
| *Lyrognathus* | *giannisposatoi* | BS |
| *Monocentropus* | *balfouri* | BS |
| *Neoholothele* | *fasciaauranigra* | BS |
| *Nhandu* | *chromatus* | BS |
| *Orphnaecus* | *sp.*8 Philippines | BS |
| *Pamphobeteus* | *antinous* | BS |
| *Pelinobius* | *muticus* | BS |
| *Phlogiellus* | *johnraylazoi* | BS |
| *Phormictopus* | *sp.*4 Cuba #1 and sp.4 Cuba #2 | BS |
| *Poecilotheria* | *metallica* and *regalis* | BS, and imgrum.pw/media/ 1879868038092146134 |
| *Psalmopoeus* | *victori* | Mendoza 2014, Figure 30 |
| *Pterinochilus* | *murinus* | Fearnottarantulas.com (chosen for observable underside) |
| *Selenocosmia* | *aruana* | BS |
| *Stromatopelma* | *calceatum* | BS |
| *Tapinauchenius* | *polybote* and *plumpies* | BS |
| *Thrigmopoeus* | *truculentus* | BS |
| *Trichopelma* | *sp.* Neiba, Dominican Rep. | BS |
| *Xenesthis* | *sp.* Colombia (poss. *immanis*) | BS |
| *Ybyrapora* | *diverspes and sooretama* | BS |

**Table S3:** Color measurements from digital photographs used for this study. 0 and 1 respectively indicate absence and presence. In the blue and green columns, * indicates genera where our scoring differs from Hsiung *et al*. (2015).

| TAXON | Blue | Green | Strid | UBris | Arb | Apo | ΔE_Lab_blue | ΔE_b_blue | ΔE_Lab_green | ΔE_a_green |
|---|---|---|---|---|---|---|---|---|---|---|
| *Acanthoscurria* | 0* | 0 | 1 | 1 | 0 | 1 | 62.20 | 65.16 | 60.23 | 43.03 |
| *Aphonopelma* | 1 | 1* | 0 | 1 | 0 | 1 | 32.05 | 26.83 | 25.52 | 15.43 |
| *Avicularia* | 1 | 1 | 0 | 1 | 1 | 1 | 36.89 | 31.25 | 32.02 | 19.44 |
| *Brachypelma* | 0 | 0 | 1 | 1 | 0 | 1 | 69.08 | 66.63 | 60.35 | 44.8 |
| *Bumba* | 1 | 0 | 1 | 1 | 0 | 1 | 44.43 | 37.96 | 76.32 | 40.75 |
| *Caribena* | 1 | 1 | 0 | 1 | 1 | 1 | 29.74 | 20.44 | 40.18 | 20.47 |
| *Catumiri* | 0 | 0 | 0 | 0 | 0 | 0 | 62.87 | 61.54 | 67.42 | 50.72 |
| *Chilobrachys* | 1 | 0 | 1 | 0 | 0 | 1 | 38.36 | 31.55 | 77.78 | 40.65 |
| *Coremiocnemis* | 1 | 0 | 1 | 0 | 0 | 1 | 43.15 | 40.02 | 80.6 | 49.39 |
| *Cyriopagopus* | 1 | 0 | 1 | 0 | 1 | 1 | 48.02 | 35.75 | 67.24 | 35.57 |
| *Cyrtopholis* | 1* | 0 | 1 | 1 | 0 | 1 | 43.85 | 36.25 | 70.93 | 39.93 |
| *Damarchus* | 1 | 0 | 0 | 0 | 0 | 0 | 46.42 | 28.49 | 72.45 | 33.89 |
| *Ephebopus* | 1 | 1 | 0 | 1 | 0 | 1 | 29.77 | 24.03 | 25.44 | 22.35 |
| *Haplocosmia* | 0* | 0 | 1 | 0 | 0 | 1 | 61.68 | 57.37 | 57.88 | 44.0 |
| *Haplopelma* | 1* | 0 | 1 | 0 | 0 | 1 | 0 | 0 | 114.72 | 62.46 |
| *Harpactira* | 1 | 1* | 1 | 0 | 0 | 1 | 28.52 | 31.77 | 47.07 | 35.04 |
| *Heteroscodra* | 0 | 0 | 0 | 0 | 1 | 0 | 70.23 | 59.36 | 53.42 | 41.69 |
| *Hysterocrates* | 0 | 0 | 1 | 0 | 0 | 1 | 65.88 | 63.85 | 60.13 | 46.27 |
| *Lasiodora* | 0 | 0 | 1 | 1 | 0 | 1 | 66.76 | 61.96 | 61.49 | 39.42 |
| *Linothele* | 1 | 0 | 0 | 0 | 0 | 0 | 44.66 | 39.48 | 72.80 | 41.6 |
| *Lyrognathus* | 0* | 0 | 1 | 0 | 0 | 1 | 57.12 | 55.0 | 67.88 | 47.56 |
| *Monocentropus* | 1 | 0 | 1 | 0 | 0 | 1 | 35.01 | 25.09 | 80.45 | 41.84 |
| *Neoholothele* | 0 | 0 | 0 | 0 | 0 | 0 | 67.72 | 65.06 | 56.26 | 45.53 |
| *Nhandu* | 0* | 0* | 1 | 1 | 0 | 1 | 61.5 | 58.12 | 59.81 | 42.51 |
| *Orphnaecus* | 1 | 0 | 1 | 0 | 0 | 1 | 31.96 | 27.21 | 84.72 | 45.81 |
| *Pamphobeteus* | 1 | 0 | 1 | 1 | 0 | 1 | 15.28 | 12.86 | 107.33 | 62.98 |
| *Pelinobius* | 0 | 0 | 1 | 0 | 0 | 1 | 65.88 | 63.75 | 60.7 | 45.89 |
| *Phlogiellus* | 1 | 0 | 1 | 0 | 0 | 1 | 39.95 | 34.11 | 76.04 | 41.71 |
| *Phormictopus* | 1 | 1* | 1 | 1 | 0 | 1 | 32.6 | 27.55 | 42.25 | 32.28 |
| *Poecilotheria* | 1 | 1 | 1 | 0 | 1 | 1 | 15.72 | 11.48 | 35.36 | 0.09 |
| *Psalmopoeus* | 1 | 0* | 1 | 0 | 1 | 1 | 66.34 | 43.07 | 55.12 | 23.82 |
| *Pterinochilus* | 1* | 1* | 1 | 0 | 0 | 1 | 45.62 | 29.05 | 29.94 | 9.14 |
| *Selenocosmia* | 0* | 0 | 1 | 0 | 0 | 1 | 57.35 | 54.36 | 62.58 | 45.52 |
| *Stromatopelma* | 0 | 1 | 0 | 0 | 1 | 0 | 84.37 | 73.71 | 43.32 | 38.5 |
| *Tapinauchenius* | 1 | 1 | 0 | 0 | 1 | 0 | 31.90 | 28.03 | 46.18 | 35.26 |
| *Thrigmopoeus* | 0* | 0 | 1 | 0 | 0 | 1 | 65.44 | 60.14 | 54.09 | 36.96 |
| *Trichopelma* | 1 | 0 | 0 | 0 | 0 | 0 | 53.52 | 46.81 | 62.11 | 37.77 |
| *Xenesthis* | 1 | 0 | 0 | 1 | 0 | 1 | 20.13 | 16.11 | 102.19 | 56.76 |
| *Ybyrapora* | 1 | 1 | 0 | 1 | 1 | 1 | 34.45 | 29.56 | 49.0 | 26.35 |

**Table S4:** Transcriptomes used in this study with information regarding species, taxonomy, tissue type of each sample (T, WB = whole body, L = legs), and respective Sequence Read Archive (SRA) accession numbers. References: B = Bond *et al*. 2014; F = Foley *et al*. 2019; S = Sanggaard *et al*. 2014.

| SPECIES | FAMILY | SUBFAMILY | T | SAMPLE ID | SRA RUN |
|---|---|---|---|---|---|
| *Acanthoscurria geniculata* | Theraphosidae | Theraphosinae | WB | SAMN 02378633 | SRR 1024075, S |
| *Aphonopelma johnnycashi* | Theraphosidae | Theraphosinae | WB | SAMN 02836947 | SRR 1514871, B |
| *Bumba cabocla* | Theraphosidae | Theraphosinae | WB | SAMN 11475839 | SRR 8944285, F |
| *Caribena versicolor* | Theraphosidae | Aviculariinae | WB | SAMN 11475841 | SRR 8944282, F |
| *Catumiri* sp. | Theraphosidae | Ischnocolinae | WB | SAMN 11475842 | SRR 8944288, F |
| *Cyriopagopus / Omothymus schioedtei* | Theraphosidae | Ornithoctoninae | WB | SAMN 11475843 | SRR 8944271, F |
| *Cyriopagopus / Omothymus* sp. "Hati Hati" | Theraphosidae | Ornithoctoninae | WB | SAMN 11475844 | SRR 8944281, F |
| *Damarchus* sp. | Nemesiidae | N/A | WB | SAMN 04453350 | SRR 3144092, B |
| *Ephebopus cyanognathus* | Theraphosidae | Psalmopoeinae | WB | SAMN 11475845 | SRR 8944275, F |
| *Haplocosmia* sp. | Theraphosidae | Selenocosmiinae | WB | SAMN 11475846 | SRR 8944280, F |
| *Haplopelma lividum / Cyriopagopus lividus* | Theraphosidae | Ornithoctoninae | L | SAMN 11475847 | SRR 8944277, F |
| *Heteroscodra maculata* | Theraphosidae | Stromatopelminae | L | SAMN 11475848 | SRR 8944261, F |
| *Hysterocrates* sp. | Theraphosidae | Eumenophorinae | L | SAMN 11475849 | SRR 8944262, F |
| *Lasiodora parahybana* | Theraphosidae | Theraphosinae | L | SAMN 11475850 | SRR 8944263, F |
| *Linothele* sp. | Dipluridae | N/A | L | SAMN 11475851 | SRR 8944264, F |
| *Monocentropus balfouri* | Theraphosidae | Eumenophorinae | WB | SAMN 11475852 | SRR 8944273, F |
| *Neoholothele incei* | Theraphosidae | Schismatothelinae | WB | SAMN 11475853 | SRR 8944287, F |
| *Pelinobius muticus* | Theraphosidae | Eumenophorinae | WB | SAMN 11475855 | SRR 8944274, F |
| *Phlogiellus inermis* | Theraphosidae | Selenocosmiinae | WB | SAMN 11475856 | SRR 8944272, F |
| *Phormictopus atrichomatus* | Theraphosidae | Theraphosinae | WB | SAMN 11475857 | SRR 8944270, F |

| | | | | | |
|---|---|---|---|---|---|
| *Poecilotheria vittata* | Theraphosidae | Poecilotheriinae | WB | SAMN 11475858 | SRR 8944269, F |
| *Psalmopoeus cambridgei* | Theraphosidae | Psalmopoeinae | WB | SAMN 11475860 | SRR 8944279, F |
| *Pterinochilus chordatus* | Theraphosidae | Harpactirinae | L | SAMN 11475862 | SRR 8944268, F |
| *Pterinochilus lugardi* | Theraphosidae | Harpactirinae | WB | SAMN 11475863 | SRR 8944284, F |
| *Selenocosmia javanensis* | Theraphosidae | Selenocosmiinae | L | SAMN 11475864 | SRR 8944260, F |
| *Stromatopelma calceatum* | Theraphosidae | Stromatopelminae | WB | SAMN 11475865 | SRR 8944276, F |
| *Tapinauchenius violaceus* | Theraphosidae | Psalmopoeinae | WB | SAMN 11475867 | SRR 8944283, F |
| *Thrigmopoeus* sp. | Theraphosidae | Thrigmopoeinae | WB | SAMN 11475866 | SRR 8944286, F |
| *Trichopelma laselva* | Theraphosidae | Ischnocolinae “sensu stricto” | WB | SAMN 02837052 | SRR 1514881, B |

**Table S5:** Reference spider opsins used in this study with respective identification numbers. Opsins were obtained per Morehouse *et al*., 2017.

| **SPECIES** | **OPSIN TYPE** | **ACCESSION NOS.** | **REFERENCE** |
|---|---|---|---|
| *Aliatypus coylei* | RH2B | PRJNA254752, SRX652492 | Bond *et al.* 2014 |
| *Aphonopelma johnnycashi* | RH2A | PRJNA254752, SRX652487 | Bond *et al.* 2014 |
| *Aphonopelma johnnycashi* | SWS | TR3076 | Morehouse *et al.* 2017 |
| *Aptostichus stephencolberti* | MWS | PRJNA254752, SRX652490 | Bond *et al.* 2014 |
| *Cuppienius salei* | RH1 | CCO61973 | Zopf *et al.* 2013 |
| *Cuppienius salei* | PER | CCP46949 | Eriksson *et al.* 2013 |

**Table S6:** Opsin orthologs and homologs across all taxa. H refers to the number of sequences that show homology to the given reference opsin, excluding predicted orthologs. O refers to the number of orthologs predicted from that species. Leg-only transcriptomes denoted by ✝.

| TAXON | SWS | | MWS | | RH2A | | RH2B | | RH1 | | RRH | |
|---|---|---|---|---|---|---|---|---|---|---|---|---|
| | H | O | H | O | H | O | H | O | H | O | H | O |
| *Acanthoscurria geniculata* | 2 | - | 2 | - | 2 | - | 2 | - | 2 | - | 2 | - |
| *Aphonopelma johnnycashi* | 2 | - | 3 | - | 2 | - | 3 | - | 3 | - | 2 | - |
| *Bumba cabocla* | - | 1 | 6 | 1 | 6 | 1 | 6 | 1 | 7 | - | 7 | - |
| *Caribena versicolor* | - | 1 | - | 1 | 1 | 1 | - | 1 | 1 | - | 2 | - |
| *Catumiri* sp. | - | 1 | 3 | 1 | 3 | 1 | 3 | 1 | 4 | - | 4 | - |
| *Cyriopagopus / Omothymus schioedtei* | 11 | - | 6 | 1 | 10 | - | 8 | - | 7 | - | 7 | - |
| *Cyriopagopus / Omothymus* sp. "Hati Hati" | - | 1 | - | 1 | 1 | 1 | 1 | 1 | 2 | - | - | 1 |
| *Damarchus* sp. | 4 | 1 | 2 | 2 | 5 | - | 6 | - | 4 | - | 1 | 1 |
| *Ephebopus cyanognathus* | 1 | - | 1 | 1 | 3 | - | 2 | 1 | 3 | - | 3 | - |
| *Haplocosmia* sp. | - | 1 | 2 | - | - | 1 | - | 1 | 1 | - | 1 | - |
| *Haplopelma lividum / Cyriopagopus lividus* ✝ | 1 | 1 | - | - | - | 1 | - | - | - | - | - | - |
| *Heteroscodra maculata* ✝ | - | 1 | - | - | - | 1 | - | 1 | - | - | - | - |
| *Hysterocrates* sp. ✝ | - | - | 2 | - | 2 | - | 2 | - | 2 | - | 5 | - |
| *Lasiodora parahybana* ✝ | - | - | - | - | - | - | - | 1 | - | - | - | - |
| *Linothele* sp. ✝ | - | - | 1 | - | 2 | 1 | 1 | - | - | - | - | - |
| *Monocentropus balfouri* | - | 1 | - | 1 | 3 | 1 | 1 | 1 | 2 | - | 1 | - |
| *Neoholothele incei* | - | 1 | - | 1 | 2 | 1 | 1 | 1 | 2 | - | 2 | - |
| *Pelinobius muticus* | 7 | 1 | 7 | 1 | 6 | 1 | 7 | 1 | 8 | - | 1 | 1 |
| *Phlogiellus inermis* | 4 | - | 5 | - | 6 | - | 4 | - | 2 | 1 | 3 | - |
| *Phormictopus atrichomatus* | - | 1 | - | 1 | 4 | 1 | 4 | 1 | 4 | - | 1 | - |
| *Poecilotheria vittata* | - | 1 | - | 1 | 2 | 1 | - | 1 | 1 | - | 1 | - |
| *Psalmopoeus cambridgei* | - | 1 | - | 1 | 5 | 1 | 5 | 1 | 6 | - | - | 1 |
| *Pterinochilus chordatus* ✝ | - | - | - | - | 2 | - | 2 | - | 2 | - | 3 | - |
| *Pterinochilus lugardi* | 1 | 1 | - | 1 | 3 | 1 | 1 | 1 | 2 | - | - | 1 |
| *Selenocosmia javanensis* ✝ | 3 | - | 2 | - | 4 | - | 3 | - | 3 | - | 7 | - |
| *Stromatopelma calceatum* | - | 2 | - | 1 | 4 | 1 | 4 | 1 | 5 | - | 5 | - |
| *Tapinauchenius voilaceus* | - | 1 | - | 1 | 2 | 1 | 2 | 1 | 3 | - | 3 | - |
| *Thrigmopoeus sp.* | - | 2 | 3 | - | 3 | 2 | 3 | - | 2 | - | 2 | - |
| *Trichopelma laselva* | - | 1 | - | 1 | 2 | 1 | 2 | 1 | - | - | 1 | - |

**Table S7:** A list of genera in this study, with the numbers of described species per genus, as of article submission date. We quantify how many unique species in each genus had useable pictures in the photo archive www.birdspiders.com and/or www.tarantupedia.com, and how many of those species were blue or green (scored visually by S. Foley). A * indicates currently undescribed species with blue or green colour that can be confidently attributed to the corresponding genus. However, as the described species exhibit no blue or green colour, we conservatively scored these as 0. For *Poecilotheria*, all species were considered both blue and green, as all known species possess iridescent green footpads.

| GENUS | # SPECIES IN GENUS | # SPECIES WITH PICTURES | NUMBER OF BLUE SPECIES | NUMBER OF GREEN SPECIES |
|---|---|---|---|---|
| *Acanthoscurria* | 27 | 12 | 0 | 0 |
| *Aphonopelma* | 59 | 37 | 5 | 2 |
| *Avicularia* | 12 | 12 | 5 | 2 |
| *Brachypelma* | 8 | 8 | 0 | 0 |
| *Bumba* | 3 | 2 | 1 | 0 |
| *Caribena* | 2 | 2 | 2 | 2 |
| *Catumiri* | 4 | 3 | 0 | 0 |
| *Chilobrachys* | 27 | 9 | 4 | 0 |
| *Coremiocnemis* | 6 | 5 | 1 | 0 |
| *Cyriopagopus* | 4 | 2 | 2 | 0 |
| *Cyrtopholis* | 24 | 7 | 1 | 0 |
| *Damarchus* | 5 | 1 | 1 | 0 |
| *Ephebopus* | 5 | 5 | 1 | 1 |
| *Haplocosmia* | 2 | 2 | 0 | 0 |
| *Haplopelma* | 10 | 7 | 2 | 0 |
| *Harpactira* | 15 | 12 | 2 | 1 |
| *Heteroscodra* | 2 | 1 | 0 | 0 |
| *Hysterocrates* | 18 | 6 | 0 | 0 |
| *Lasiodora* | 33 | 5 | 0 | 0 |
| *Linothele* | 26 | 1 | 1 | 0 |
| *Lyrognathus* | 7 | 6 | 0 | 0 |
| *Monocentropus* | 3 | 1 | 1 | 0 |
| *Neoholothele* | 2 | 2 | 0 | 0 |
| *Nhandu* | 5 | 4 | 0 | 0 |
| *Orphnaecus* | 4 | 3 | 0* | 0 |
| *Pamphobeteus* | 13 | 6 | 4 | 0 |
| *Pelinobius* | 1 | 1 | 0 | 0 |
| *Phlogiellus* | 23 | 11 | 5 | 0 |
| *Phormictopus* | 14 | 2 | 1 | 0* |
| *Poecilotheria* | 15 | 15 | 15 | 15 |
| *Psalmopoeus* | 14 | 6 | 1 | 0 |
| *Pterinochilus* | 10 | 5 | 1 | 1 |
| *Selenocosmia* | 36 | 6 | 0 | 0 |
| *Stromatopelma* | 6 | 1 | 0 | 1 |
| *Tapinauchenius* | 9 | 6 | 4 | 3 |
| *Thrigmopoeus* | 2 | 2 | 0 | 0 |
| *Trichopelma* | 17 | 0 | 0* | 0 |
| *Xenesthis* | 3 | 3 | 1 | 0 |
| *Ybyrapora* | 3 | 3 | 3 | 2 |

**Figure S1:** Ancestral state reconstruction of blueness (ΔE *Lab*) using *ancThresh* after 10 million generations. Posterior probability densities for each ancestral node are also shown alongside.

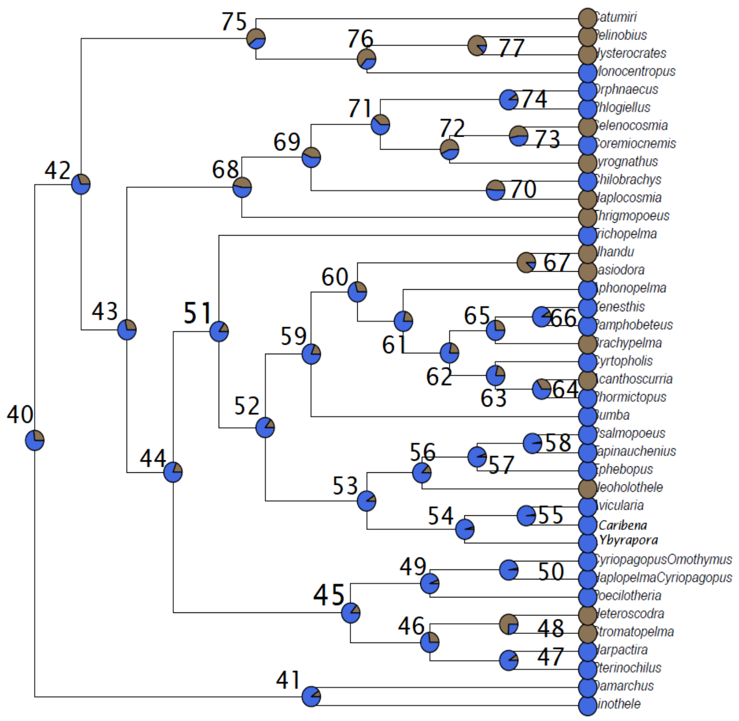


| Node# | burlywood4 | royalblue |
|---|---|---|
| 40 | 0.26959130 | 0.7304087 |
| 41 | 0.09861267 | 0.9013873 |
| 42 | 0.30296213 | 0.6970379 |
| 43 | 0.28158980 | 0.7184102 |
| 44 | 0.19335083 | 0.8066492 |
| 45 | 0.14298213 | 0.8570179 |
| 46 | 0.26771654 | 0.7322835 |
| 47 | 0.09536308 | 0.9046369 |
| 48 | 0.74365704 | 0.2563430 |
| 49 | 0.06186727 | 0.9381327 |
| 50 | 0.03162105 | 0.9683790 |
| 51 | 0.15560555 | 0.8443945 |
| 52 | 0.14798150 | 0.8520185 |
| 53 | 0.09461317 | 0.9053868 |
| 54 | 0.04161980 | 0.9583802 |
| 55 | 0.02112236 | 0.9788776 |
| 56 | 0.12985877 | 0.8701412 |
| 57 | 0.05374328 | 0.9462567 |
| 58 | 0.02712161 | 0.9728784 |
| 59 | 0.19060117 | 0.8093988 |
| 60 | 0.28946382 | 0.7105362 |
| 61 | 0.21809774 | 0.7819023 |
| 62 | 0.22047244 | 0.7795276 |
| 63 | 0.21372328 | 0.7862767 |
| 64 | 0.33858268 | 0.6614173 |
| 65 | 0.26459193 | 0.7354081 |
| 66 | 0.09186352 | 0.9081365 |
| 67 | 0.88213973 | 0.1178603 |
| 68 | 0.46494188 | 0.5350581 |
| 69 | 0.43369579 | 0.5663042 |
| 70 | 0.48268966 | 0.5173103 |
| 71 | 0.37832771 | 0.6216723 |
| 72 | 0.57180352 | 0.4281965 |
| 73 | 0.53280840 | 0.4671916 |
| 74 | 0.09373828 | 0.9062617 |
| 75 | 0.60692413 | 0.3930759 |
| 76 | 0.63804524 | 0.3619548 |
| 77 | 0.87701537 | 0.1229846 |

**Figure S2:** Ancestral state reconstruction of greenness (ΔE *Lab*) using *ancThresh* after 10 million generations. Posterior probability densities for each ancestral node are shown alongside.

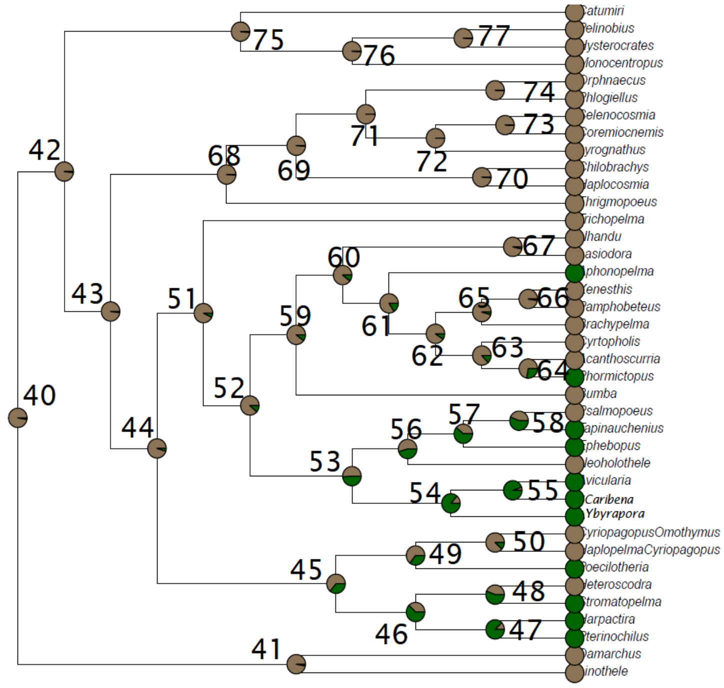


| Node# | burlywood4 | darkgreen |
|---|---|---|
| 40 | 0.9698788 | 0.030121235 |
| 41 | 0.9693788 | 0.030621172 |
| 42 | 0.9801275 | 0.019872516 |
| 43 | 0.9782527 | 0.021747282 |
| 44 | 0.9513811 | 0.048618923 |
| 45 | 0.6419198 | 0.358080240 |
| 46 | 0.3724534 | 0.627546557 |
| 47 | 0.1227347 | 0.877265342 |
| 48 | 0.4391951 | 0.560804899 |
| 49 | 0.6456693 | 0.354330709 |
| 50 | 0.8798900 | 0.120109986 |
| 51 | 0.9351331 | 0.064866892 |
| 52 | 0.8797650 | 0.120234971 |
| 53 | 0.5224347 | 0.477565304 |
| 54 | 0.1423572 | 0.857642795 |
| 55 | 0.0543682 | 0.945631796 |
| 56 | 0.5469316 | 0.453068366 |
| 57 | 0.3990751 | 0.600924884 |
| 58 | 0.4545682 | 0.545431821 |
| 59 | 0.9022622 | 0.097737783 |
| 60 | 0.8922635 | 0.107736533 |
| 61 | 0.8302712 | 0.169728784 |
| 62 | 0.9020122 | 0.097987752 |
| 63 | 0.8692663 | 0.130733658 |
| 64 | 0.7289089 | 0.271091114 |
| 65 | 0.9541307 | 0.045869266 |
| 66 | 0.9730034 | 0.026996625 |
| 67 | 0.9643795 | 0.035620547 |
| 68 | 0.9883765 | 0.011623547 |
| 69 | 0.9943757 | 0.005624297 |
| 70 | 0.9947507 | 0.005249344 |
| 71 | 0.9953756 | 0.004624422 |
| 72 | 0.9957505 | 0.004249469 |
| 73 | 0.9956255 | 0.004374453 |
| 74 | 0.9941257 | 0.005874266 |
| 75 | 0.9851269 | 0.014873141 |
| 76 | 0.9910011 | 0.008998875 |
| 77 | 0.9908761 | 0.009123860 |

**Figure S3:** Full ancestral state reconstruction of arboreality using *ancThresh* after 10 million generations. Posterior probability densities for each ancestral node are shown alongside (light blue shading corresponds to arboreality, brown to non-arboreality).

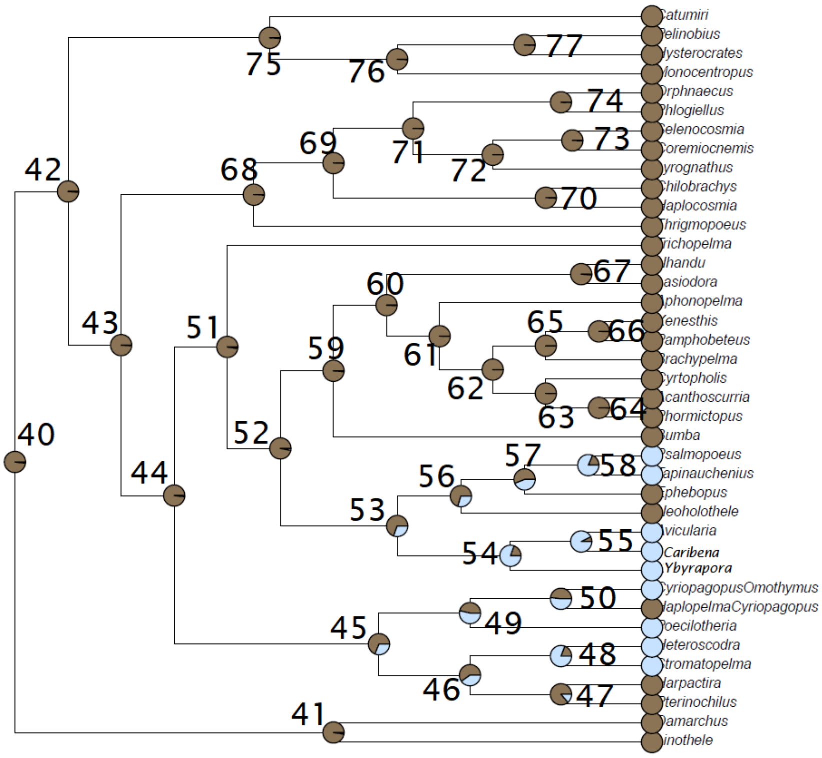


| Node# | burlywood4 | slategray1 |
|---|---|---|
| 40 | 0.98312711 | 0.016872891 |
| 41 | 0.97937758 | 0.020622422 |
| 42 | 0.98900137 | 0.010998625 |
| 43 | 0.99050119 | 0.009498813 |
| 44 | 0.98225222 | 0.017747782 |
| 45 | 0.68378953 | 0.316210474 |
| 46 | 0.60104987 | 0.398950131 |
| 47 | 0.86689164 | 0.133108361 |
| 48 | 0.19735033 | 0.802649669 |
| 49 | 0.46681665 | 0.533183352 |
| 50 | 0.47781527 | 0.522184727 |
| 51 | 0.97712786 | 0.022872141 |
| 52 | 0.97400325 | 0.025996750 |
| 53 | 0.70116235 | 0.298837645 |
| 54 | 0.19835021 | 0.801649794 |
| 55 | 0.07611549 | 0.923884514 |
| 56 | 0.69728784 | 0.302712161 |
| 57 | 0.56180477 | 0.438195226 |
| 58 | 0.19010124 | 0.809898763 |
| 59 | 0.98312711 | 0.016872891 |
| 60 | 0.99187602 | 0.008123985 |
| 61 | 0.99450069 | 0.005499313 |
| 62 | 0.99750031 | 0.002499688 |
| 63 | 0.99662542 | 0.003374578 |
| 64 | 0.99662542 | 0.003374578 |
| 65 | 0.99800025 | 0.001999750 |
| 66 | 0.99762530 | 0.002374703 |
| 67 | 0.99000125 | 0.009998750 |
| 68 | 0.99350081 | 0.006499188 |
| 69 | 0.99750031 | 0.002499688 |
| 70 | 0.99462567 | 0.005374328 |
| 71 | 0.99687539 | 0.003124609 |
| 72 | 0.99712536 | 0.002874641 |
| 73 | 0.99737533 | 0.002624672 |
| 74 | 0.99462567 | 0.005374328 |
| 75 | 0.98900137 | 0.010998625 |
| 76 | 0.99162605 | 0.008373953 |
| 77 | 0.99137608 | 0.008623922 |

**Figure S4:** The pattern of opsin evolution in tarantulas. The long-wavelength RH2B opsin clade emerges as the most basal, followed by RH2A, medium-wave (MWS), short-wave (SWS), with peropsins (RRH) as the most derived. While this pattern is largely consistent with the literature (e.g. Biscontin et al., 2016), it differs in that peropsins do not emerge as the most basal in our case.

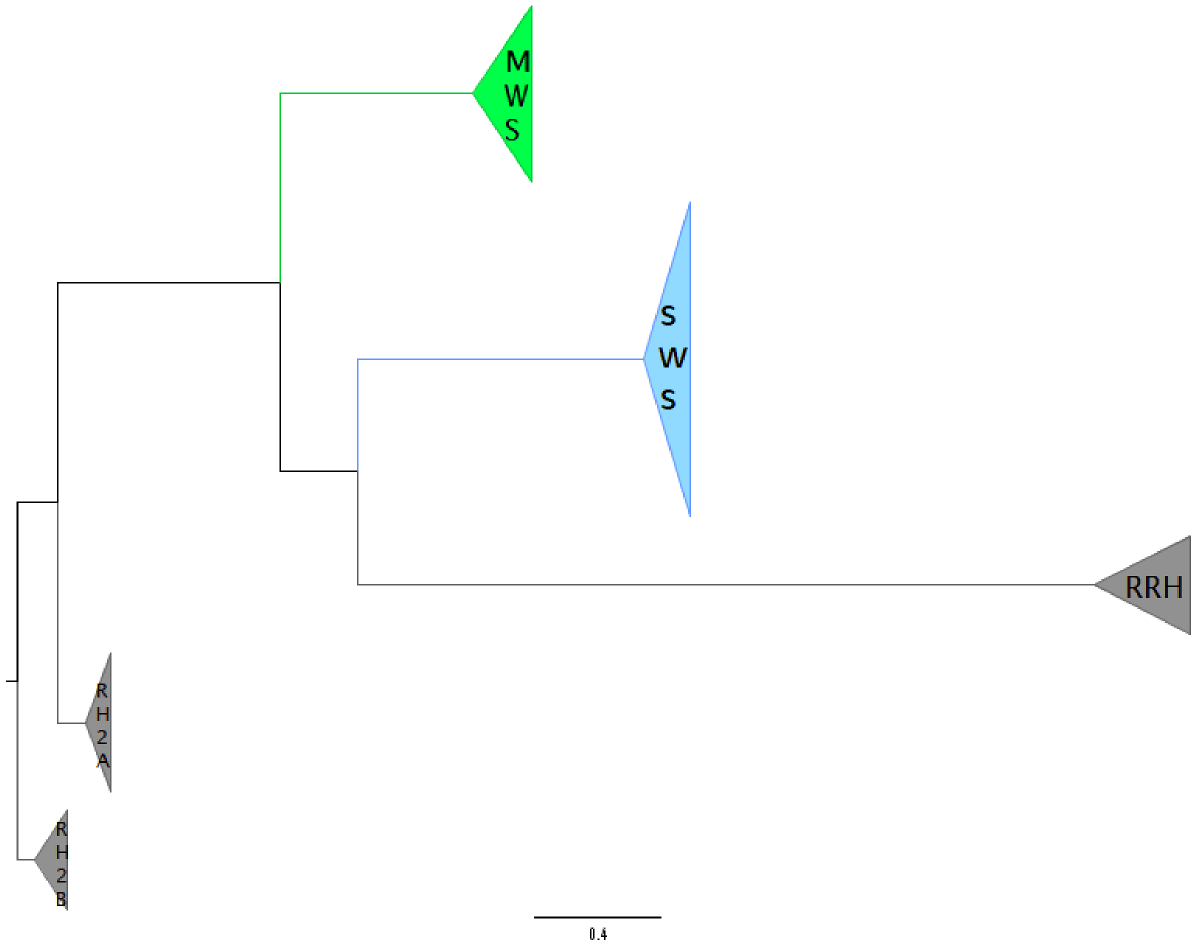

**Figure S5:** Multiple Sequence Alignments of SWS opsins (top), MWS opsins (second), RH2A (third), RH2B (fourth), and RRH (bottom). Reference sequences are the top-most in each case.

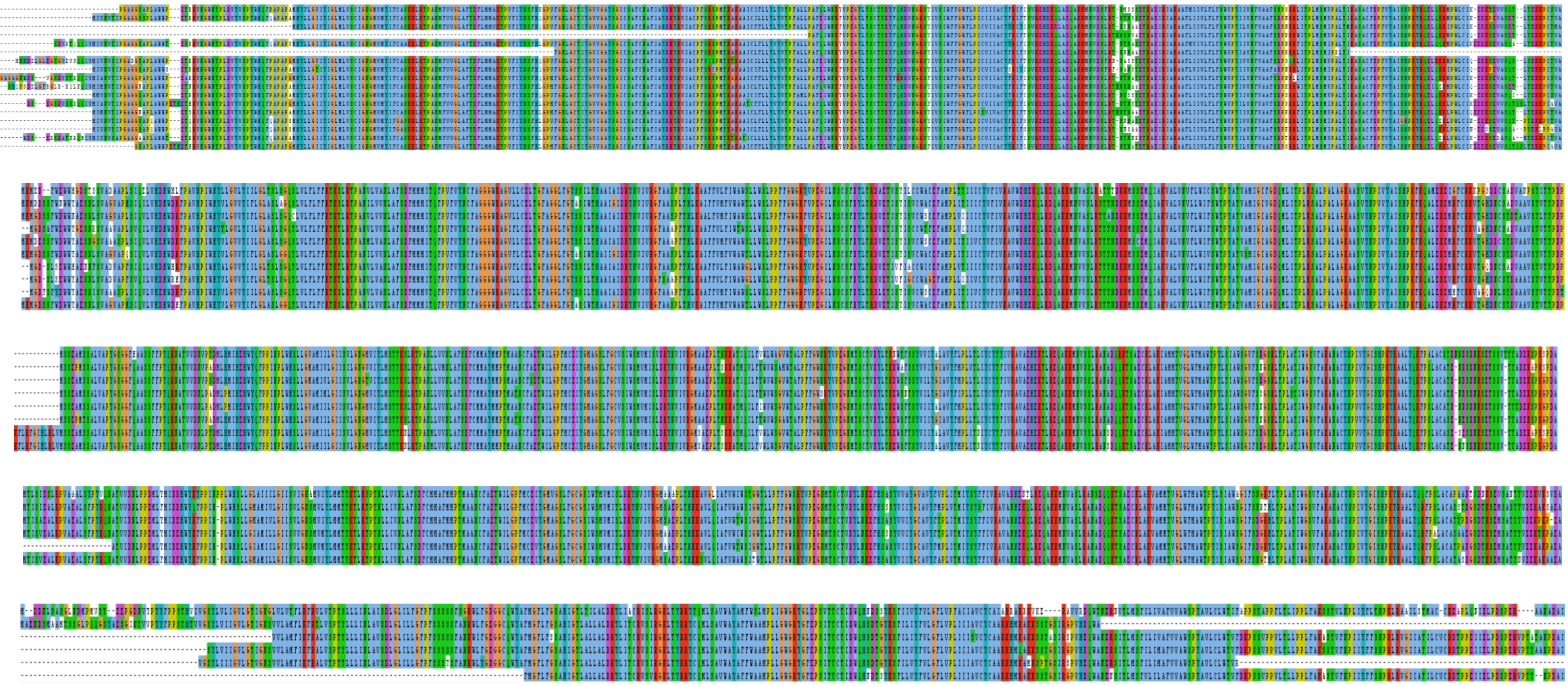
